# Highly parallelized construction of DNA from low-cost oligonucleotide mixtures using Data-optimized Assembly Design and Golden Gate

**DOI:** 10.1101/2023.11.20.567888

**Authors:** Sean Lund, Vladimir Potapov, Sean R. Johnson, Jackson Buss, Nathan A. Tanner

**Affiliations:** Research Department, New England Biolabs, Ipswich, MA 01938, USA

**Keywords:** DNA synthesis, Golden Gate Assembly

## Abstract

Commercially synthesized genes are typically made using variations of homology-based cloning techniques, including polymerase cycling assembly from chemically synthesized microarray-derived oligonucleotides. Here we apply Data-optimized Assembly Design to the synthesis of hundreds of codon-optimized genes in both constitutive and inducible vectors using Golden Gate Assembly. Starting from oligonucleotide pools, we synthesize genes in three simple steps: 1) Amplification of parts belonging to individual assemblies in parallel from a single pool; 2) Golden Gate Assembly of parts for each construct; and 3) Transformation. We construct genes from receiving DNA to sequence confirmed isolates in as little as 4 days. By leveraging the ligation fidelity afforded by T4 DNA ligase, we expect to be able to construct a larger breadth of sequences not currently supported by homology-based methods which require stability of extensive single-stranded DNA overhangs.

## 1. Introduction

DNA construction is central to testing biological hypotheses or engineering using the Design-Build-Test-Learn cycle. While DNA costs have lowered, they have stagnated^1^. DNA construction is primarily a service controlled by and obtained from DNA providers^2^. We propose an improvement on gene construction synthesis schemes using oligonucleotide (oligo) pools as sources of user-defined DNA sequences for decentralized gene construction through Golden Gate Assembly following a simple three-step process.

Complex DNA synthesis has historically been accomplished by synthesizing overlapping single-stranded DNA (ssDNA) from microarray or column synthesis, annealing, and generation of double-stranded DNA (dsDNA) fragments by polymerase-cycling assembly^3-6^. Genes are constructed in parallel or in some instances, a pooled manner^7^. In order to use low-cost microarray-derived DNA, this process required tedious and lengthy procedures to first fish out oligos from complex mixtures followed by generation of ssDNA through treatments like the use of Lambda exonuclease to preferentially digest phosphorylated strands in dsDNA duplexes^4^. This procedure has been modified to utilize dsDNA fragments directly by removing barcodes using Type IIS restriction enzymes followed by generation of short segments of 3’ overhangs with T5 exonuclease in Gibson Assembly^8^. Unfortunately, both procedures generate extensive segments of ssDNA potentially prone to forming secondary structures or serving as substrates for single-stranded endonuclease activity of T5 exonuclease, thereby reducing the efficiency and accuracy of assembly^9^. Many of these failures can be predicted, but still limit the space of sequences that can be reliably synthesized.

In contrast to the stretches of ssDNA homology required in Gibson assembly^10^, Golden Gate Assembly utilizes short overhangs generated by restriction digestion with the same Type IIS enzymes used to remove barcodes in classical dsDNA construction schemes^4^. These short overhangs are then enzymatically ligated by the ligase in a single-pot reaction^11^. Overhang sets are often chosen by rules of thumb and used in tiered-cloning schemes limiting the number of parts that can be successfully combined^12, 13^. These overhangs can be methodically selected using ligation fidelity data in a process referred to as Data-optimized Assembly Design (DAD) which has been shown to support assemblies in as high complexity as 52 fragments^14-16^.

Here we apply DAD for gene construction from oligo pools using the fragment design NEBridge SplitSet® Tool (Figure 1). Protein sequences are first codon optimized to eliminate Type IIS restriction sites before DAD is applied to optimize the splitting of sequences into fragments that can be synthesized on a microarray. Individual designs are then barcoded and sequences from all designs are pooled and synthesized. Individual designs are then retrieved in a multiplexed PCR using the design’s assigned barcode set before being purified as a pool and then assembled into a vector. Following transformation of the Golden Gate Assembly, 4 colonies are picked for colony PCR and amplicons are pooled for sequencing on a MinION (Oxford Nanopore Technologies) instrument using a previously validated scheme^17^. The construction process can be conducted in less than one week. We believe that Golden Gate paired with DAD presents significant benefits to homology-based DNA assembly from microarray-derived oligos.

**Figure 1.**
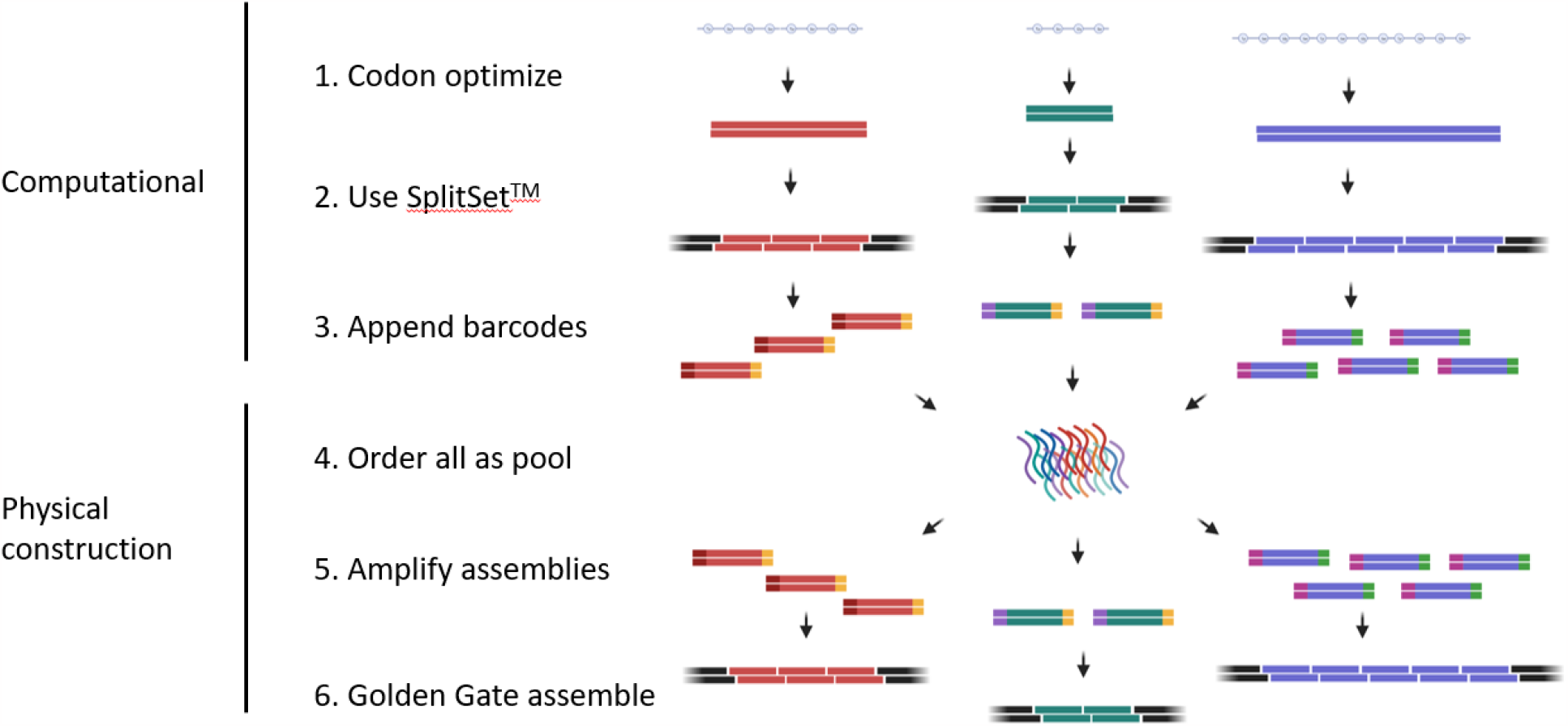
Schematic showing assembly of genes using oligo pools. Genes are first codon optimized to eliminate internal BsaI sites. Next the sequences are fragmented into pieces by using NEBridge SplitSet® Tool. The pieces are then barcoded and pooled for ordering. Individual designs are then retrieved by PCR using a design’s barcode set before purification and assembly by Golden Gate. Created with BioRender.com.

## 2. Results

### 2.1 Investigating different oligo layouts by construction of difficult sequences

Oligo layouts were expected to influence construction rates based on the number of fragments involved in an assembly as well as the ability to parallelize PCR reactions and subsequent clean-ups. Four different oligo design schemes consisting of different length oligos and stuffer arrangements were evaluated (Figure 2). Two different oligo lengths were investigated: 200 nt and 300 nt pools. To obtain oligos of equal length, oligos for the construction were padded with additional sequence to obtain oligos of the desired uniform length. This padding was either done in between the primer binding sites used for amplification or outside of the primers. If the padding is done between the primers, PCR amplicons will be of equal length while padding outside of the primers would result in pooled amplifications yielding products of various lengths.

**Figure 2.**
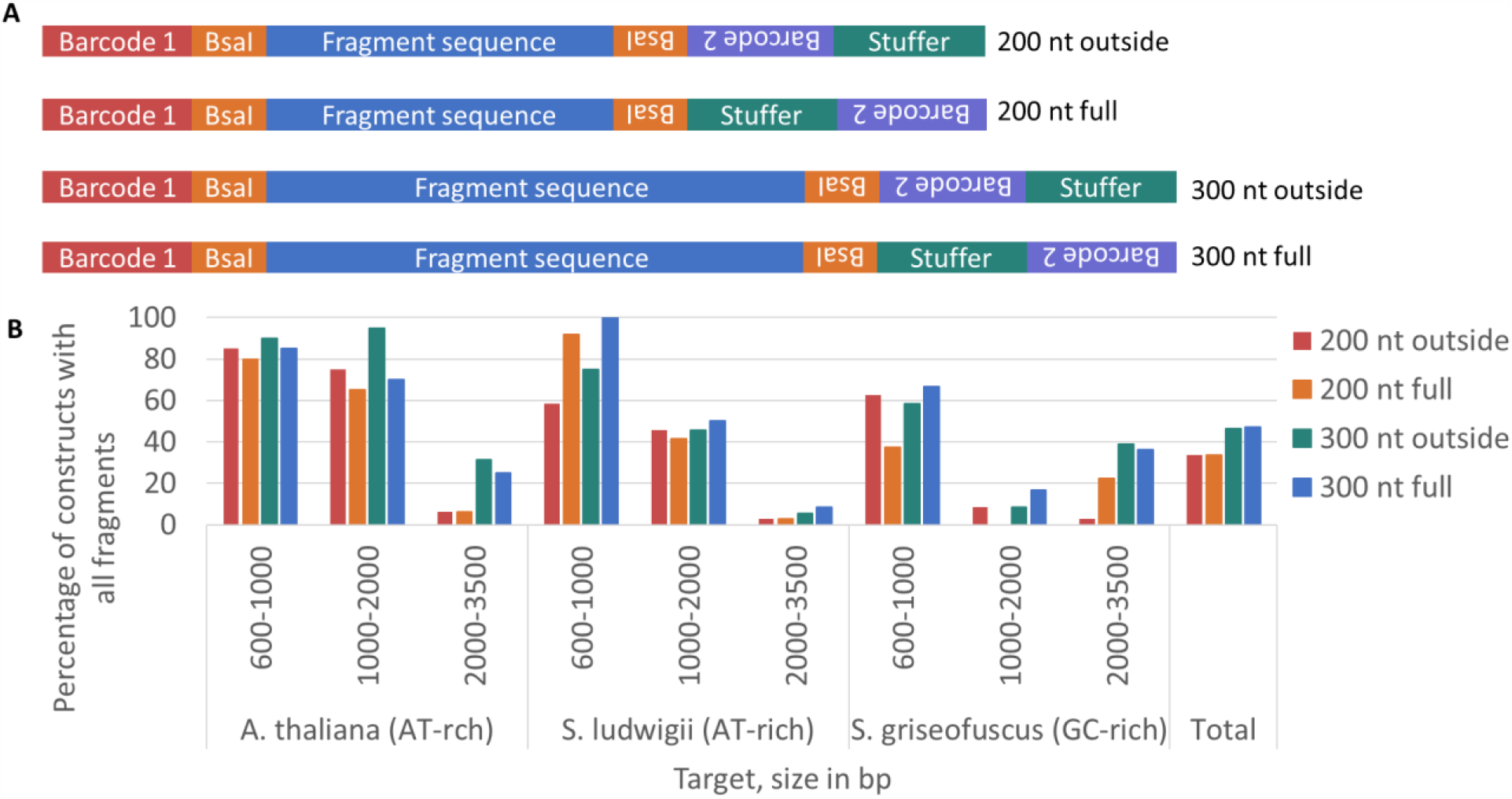
Examining different oligo layouts and their effect on success of fragment assembly. (A) Schematic depicting various oligo layouts for examination. (B) Results of attempted construction of various difficult sequences. *A. thaliana* sequences are AT-rich, *S. ludwigii* are AT-rich, and *S. griseofuscus* are GC-rich. Various length constructs were attempted for construction with longer sequences being more difficult to synthesize. Construction rates were validated on the inclusion of all fragments as MinION sequencing was unable to differentiate correct versus incorrect sequences due to the presence of homopolymers.

To test these layouts, sequences were pooled from *Arabidopsis thaliana* (AT-rich, 36% GC), *Saccharomycodes ludwigii* (AT-rich, <30% GC), or *Streptomyces griseofuscus* (GC-rich, >70% GC). 18 different assemblies of naturally occurring sequences between 600 and 3500 bp of each were attempted with the majority of the targets not eligible for dsDNA fragment synthesis by popular gene synthesis companies according to online portals (Supplementary Figure 1, Supplementary File 1), due to falling outside of acceptable specifications for GC content, repeat content, or other sequence parameters. Isolates were analyzed for their ability to incorporate every piece of the assembly rather than the sequence correctness (Figure 2). While many of these difficult to synthesize sequences were not 100% correct, several were successfully synthesized, including sequences of >70% or <30% GC content and those with indirect or direct repeats longer than 10 bases. Many of these sequences had extensive homopolymer regions or repeats incompatible with MinION sequencing limiting the analysis to which isolates contained all the desired fragments^18^.

For the 200 nt pool, we found an isolate containing all 22 pieces in the assembly and for the 300 nt pool, we found an assembly accommodating 12 pieces. While 200 nt pools supported assemblies with more pieces in certain instances, in general longer sequences could be constructed from 300 nt pools due to the longer length of each piece. For this reason, the 300 nt pools were further pursued.

In general, it was found that the 200 nt oligo pools had lower success rates for longer products than the 300 nt pools presumably due to an inverse relationship between length of oligo and the complexity of multiplexed PCR and subsequent assembly. These studies were limited to PCR amplification with Q5, obscuring the impact of choice of polymerase on the construction rate due to factors such as GC-bias. While we expected to find an impact of stuffer layout on the ability to construct target sequences, we found negligible differences between layouts in both the 200 and 300 nt pools. It was found that the AT-rich *A. thaliana* and sequences of *S. ludwigii* were constructed at higher rates than the GC-rich sequences of *S. griseofuscus*. Sequences as large as 2 kb could be built without any errors and sequences as large as 3 kb assembled but, contained sequence errors. While the stuffer layout had no apparent impact, the full-length PCRs were chosen due to the presumed benefit of the corresponding homogeneity of the multiplexed PCR for following experiments.

### 2.2 Template input requirements for single PCR retrieval

Eight designs of 9-and 8-piece assemblies from another order were selected from various problematic and successful assemblies using the bottom layout in Figure 2 in which PCR reactions generated pooled amplicons of equal length. In contrast to published methods using a series of amplifications to enrich for parts in a design^4^, we attempted to use a single amplification from the complex mixture to retrieve fragments. Using previously validated orthogonal primer sets^19^, we amplified each design at four different template concentrations. After assembly, we sequenced four isolates per design and template concentration combination (Figure 3). We found that higher amounts of template led to an increase in the percentage of isolates harboring all parts. When ample template was added, we found a corresponding decrease in the percentage of isolates missing fragments, parental vectors, and unexplained vector closure events which dominated the lowest input assemblies. A typical oligo pool from Twist Bioscience delivers roughly 100 ng. At this yield, roughly 20 96-well plates of amplification can be performed from a single pool.

**Figure 3.**
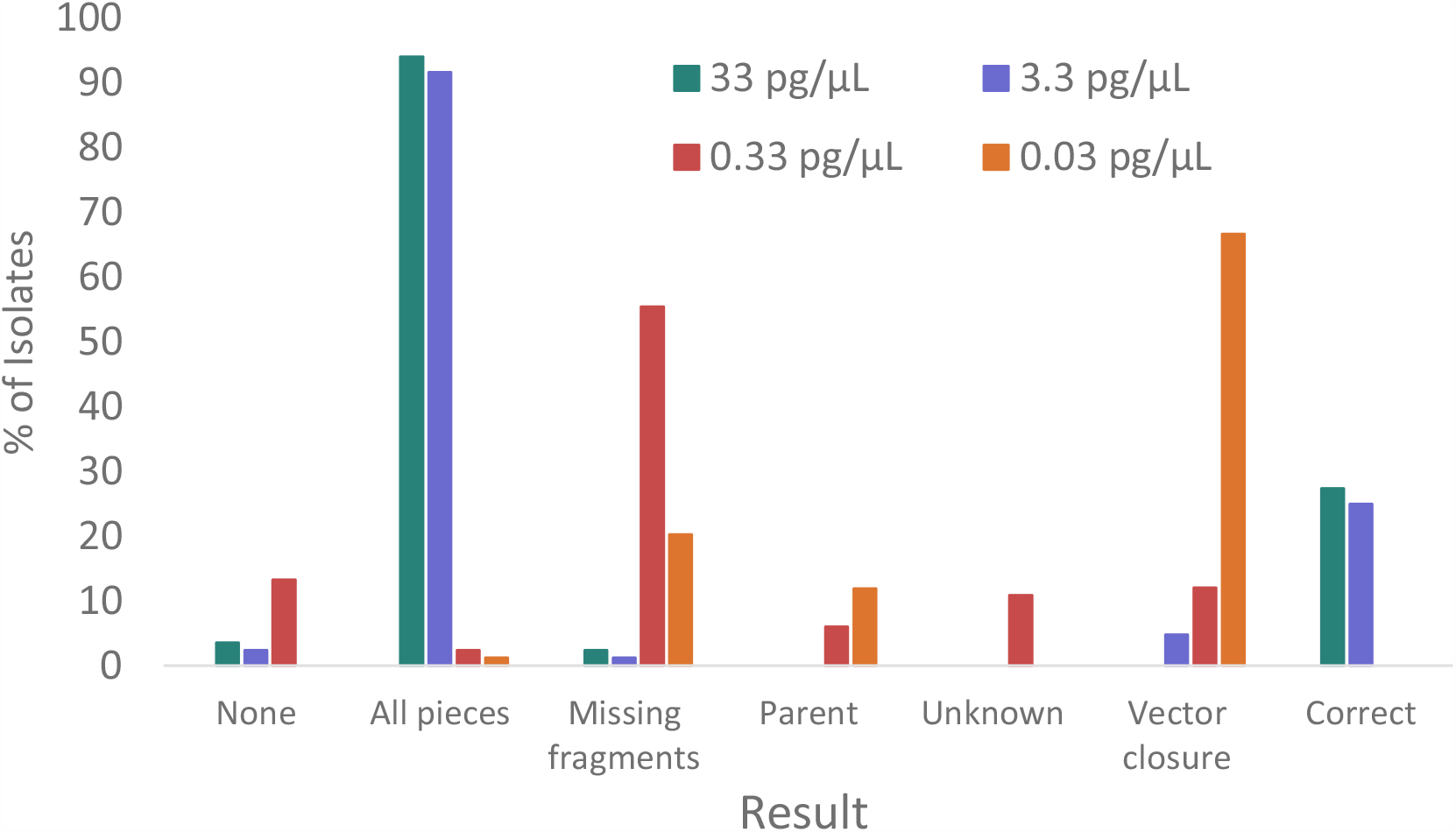
Titrating the amount of template needed for successful assembly. Various constructs were attempted for building and analyzed for the assembly result. Template concentrations greater than 3.3 pg/μL led to >90% of colonies having assemblies with all fragments present. None refers to no amplicon consensus sequenced obtained. Unknown refers to consensus sequences that could not be mapped to the parent vector. Vector closure events represent isolates where the parent vector closed without the inclusion of the original multiple cloning site flanked by BsaI sites or any intended fragments. Correct refers to isolates found that were 100% sequence correct while All pieces refers to isolates in which all fragments in the assembly could be accounted for in the isolate sequence.

### 2.3 Attempted construction of 458 genes from 2 oligo pools

To test the method in practical synthetic biology workflows, a selection of protein sequences for study were gathered from various researchers in the Research Department at New England Biolabs. These 458 proteins included enzymes hypothesized to modify DNA, protein folding chaperones, polymerases, and various phage enzymes. These enzymes included thymidylate synthase homologs expected to interfere with nucleoside metabolism, phage ten eleven translocase (TET) enzymes involved in 5-methylcytidine oxidation, and polymerases responsible for telomere maintenance in *Actinomycetota*. In addition to their questionable toxicity, some researchers requested that their proteins be cloned under constitutive promoters. In all 288 genes were attempted for cloning in a constitutive vector and the other 170 genes were cloned into a domesticated pET vector according to the desire of the researchers to whom the constructs would be delivered.

Genes were codon optimized with the restriction that they not contain BsaI sites^20^. Codon optimized genes were then run through NEBridge SplitSet® Tool to obtain fragments which were ordered across two oligo pools using the layout in which PCR generated a full 300 bp product. Three runs of 96 samples at a time, and one run of 192 samples at a time were performed. Some genes were attempted twice to deliver more constructs to researchers, but the analysis herein is limited to the first attempt and 4 colonies sequenced for each construct.

Of the 458 genes attempted for construction, 343 of the genes were successfully built without errors after checking 4 colonies (Figure 4A, Supplementary File 2). Cumulatively, greater than 80% of genes less than 12 fragments were constructed. A correlation between the number of fragments and length of construct puts the average fragment length at 220 bp (Supplementary Figure 2). Based on this, one could reasonably expect to construct sequences near 2.5 kb in length without errors, which suggests the effects of limiting by the error rate of oligo synthesis advertised by the manufacturer (1:3000 nt)^21^.

**Figure 4.**
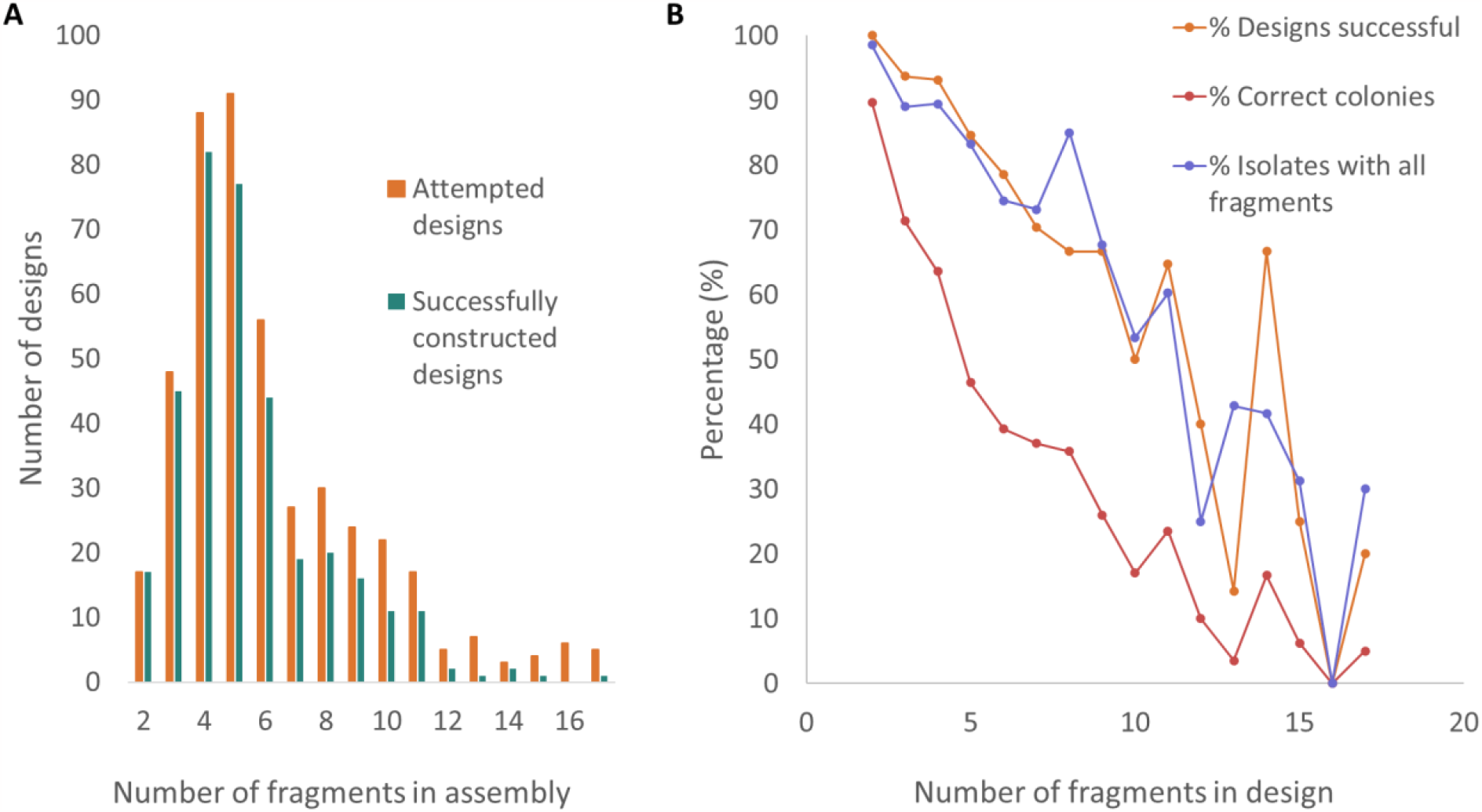
Results of attempting to construct 458 genes. (A) Overlap of design space compared to constructs successfully built after sequencing 4 colonies from each design. (B) Plot showing the relationship between fragment count and success of a given colony or design.

As the size of the assembly increases, both the number of colonies containing all fragments as well as 100% sequence correct isolates drop (Figure 4B, Supplementary Figure 3). At 12 fragments, 10% of colonies are sequence correct while for 40% of designs, a correct isolate was identified. This indicated that the success rate of a given design was likely dependent on the sequence itself, as some constructs had multiple positive isolates while others had none. As with any DNA sequence being heterologously manipulated, this could be due to the toxicity of the sequence being constructed as some of the genes being built were hypothesized to modify DNA or were constitutive in the host housing the construction. We observed some correlation between the predicted assembly efficiency and the number of colonies with all fragments (Supplementary Figure 4).

While we could not distinguish between the incorrect ligation of fragments and the errors introduced by DNA synthesis, we did observe common failure modes attributed to the nature of increasing our throughput. For some construction sets, we observed a high rate of parental vector recovery in which the multiple cloning site sequence was ligated back into the digested vector. For other sets we observed contamination of our backbone vector with a plasmid not containing any BsaI sites. When looking at vector closure events, we noticed a predominant failure mechanism by which the 3’ overhangs were presumably complemented by *in vivo* 5’ extension followed by blunt-end ligation (Supplementary Figure 5). We also observed similar events in which 5’-3’ exonuclease digestion blunted 3’ overhangs followed by some sort of blunt end ligation. We expect that these ligation events occur *in vivo* under antibiotic selection of intact vector.

## 3. Discussion

We were able to successfully convert standard homology-based gene synthesis workflows to one dependent on ligase fidelity using Golden Gate Assembly. By titrating the amount of template used in amplification of individual designs, we were able to forgo the need to amplify parts over several successive and different PCR reactions as has been used in previous schemes. While we constructed many naturally occurring AT and GC-rich sequences, reliance on sequencing chemistry limited our ability to verify sequence construction of some products and reliance on current chemistry-based synthesis methodologies limited the range of oligo sequences available as substrates. In our studies presented here, we did not distinguish between failure modes of toxicity, synthesis, sequencing, or ligation fidelity.

We successfully constructed several sequences that were rejected by various manufacturers’ web portals. If indeed these limitations are due to homology-based construction methods, Golden Gate here proves to be a valuable tool in constructing traditionally difficult to synthesize sequences. In total, 389 kb of functional sequence was successfully synthesized from a total of 615 kb of functional DNA ordered. As expected, longer sequences had higher failure rates due to both the complexity of the Golden Gate Assembly as well as the error rate in oligonucleotide synthesis. To increase the success of finding sequence correct colonies, choosing more than 4 colonies for sequencing should allow for the construction of larger sequences. We were able to sequence 762 isolates in a single run on a MinION flow cell using a barcoded cPCR scheme which could be expanded as the coverage of individual amplicons was greater than 1000.

Here we show the ability to directly amplify parts belonging to an assembly from a pool without multiple rounds of PCR, assembly using DAD and Golden Gate Assembly, and direct transformation of assemblies without an additional round of purifying PCR. With enzymatic DNA synthesis showing promise towards building sequences not accessible by chemical synthesis, we expect methods such as the ones presented here will help enable building of larger domesticated constructs^22^. As oligo pools become a more standard form of ordering novel sequences and are deployed^23^, Golden Gate Data-optimized Assembly Design will provide a compelling method in which to access low-cost DNA for construction of larger sequences.

## 4. Materials and Methods

### 4.1 Oligo pool ordering

Genes were codon optimized using DNAChisel^20^ with the restrictions of not containing any BsaI sites. To the 5’ end, <u>GGTCTCG</u>A was added such that digestion resulted in a AATG overhang. To the 3’ end, ATTC<u>CGAGACC</u> was added such that digestion resulted in an ATTC overhang on the 3’ end of the vector. Genes were then run through NEBridge SplitSet® Tool to split genes into optimized Golden Gate Assembly fragments using a faster optimization algorithm to allow for processing many constructs. The fragments were then padded with additional non-coding sequence to make the fragments approximately 260 nt in length (depending on the result from NEBridge SplitSet® Tool) before barcodes were appended to the 5’ and 3’ end^19^. The resulting scheme ensured that each oligo from a design resulted in a product 300 nt in length with the same combination of primer binding sites at each end. This was repeated for all designs with each design assigned its own unique primer barcode combination drawn from a total of 96 barcodes screened to not contain BsaI sites (Supplementary Table 1). Designed sequences were then ordered as oligonucleotide pools (Twist Biosciences, South San Franscisco, CA USA).

### 4.2 Gene construction from oligo pools

Acceptor vectors were constructed such that insertions of codon optimized genes all utilized the same 5’ and 3’ terminal overhangs AATG and ATTC, respectively using NEBuilder® HiFi DNA Assembly from templates linearized by PCR and dsDNA fragments (gBlocks, Integrated DNA Technologies, Coralville, IA USA) (Supplementary Files 3 and 4). Upstream of the promoter and downstream of the terminator primer, binding sites were added for insert sequencing. Two accepting vectors were constructed from BsaI-free domesticated backbones: pET vector based on a pET28a backbone utilizing a T7 promoter and a pAL vector based on a pACYC vector but with a rrnB P1 constitutive promoter and a series of rrnB terminators.

Oligonucleotide pools were resuspended to 1 ng/μL. A master mix of Q5® Hot Start High-Fidelity 2X Master Mix (M0494, NEB) containing oligo pool template at 3.33 pg/μL and LAMP Fluorescent Dye (B1700, NEB) at 0.5X was aliquoted across a PCR plate. Primers were then cherry-picked from a 384PP plate containing primers at 100 μM using an Echo 525 to a final concentration of 0.5 μM. A PCR was then run using an initial denaturation of 98 °C for 30 seconds before cycling 98 °C for 5 seconds, 64 °C for 30 seconds, and 72 °C for 30 seconds on a Bio-Rad CFX96 qPCR Real-Time instrument. Cycles were stopped before all amplifications plateaued in fluorescence. PCRs were then purified using 1.8X SPRISelect beads (Beckman Coulter, Brea, CA USA). Beads were washed twice using freshly prepared 80% ethanol before being eluted in nuclease free water in a volume equivalent to the initial PCR volume. Purified PCR concentrations were then spot-checked yielding between 10 and 40 ng/μL using a NanoDrop™ One Microvolume UV-Vis Spectrophotometer (ThermoFisher Scientific, Waltham, MA USA).

A master mix (2.5 μL total) of acceptor vector (13-26 fmol), 0.5 μL BsaI-HF® v2 NEBridge® Golden Gate Assembly mix (E1601, NEB, 5 μL final volume), and 0.5 μL T4 DNA ligase buffer was aliquoted across a PCR plate. 2.5 μL of purified PCR product was added and the reaction mixture mixed. The reaction was then cycled 60 times for 37 °C for 5 minutes followed by 16 °C for 5 minutes. The final reaction was held at 55 °C for 15 minutes before resting at 12 °C until transformation.

2 μL of each assembly was then transformed in parallel into NEB® 10-beta Competent *E. coli* (C3019P, NEB). Mixtures were allowed to incubate on ice for 30 minutes before a 30 second heat shock at 42 °C on a thermocycler using the lid set to the same temperature. The mixture was then returned to ice for 5 minutes before 180 μL of NEB® 10-beta/Stable Outgrowth Medium (B9035, NEB) was added. The mixture was incubated on a thermocycler at 37 °C for 1.5 hours. The mixture was then serially diluted 5-fold 4 times and the dilutions plated on Nunc™ OmniTray™ LB agar plates supplemented with the appropriate antibiotic. Plates were seeded by gently spotting 7 μL of dilutions onto plates wherein each droplet was spatially separated in a 96-well plate array using a Viaflo 96 (Integra Biosciences, Hudson, NH USA). Plates were subsequently grown at 30 °C overnight.

### 4.3 Isolate sequencing confirmation

96 unique barcodes were each appended to primers annealing upstream of the constructed operon. 16 unique barcodes were each appended to primers annealing downstream of the constructed operon. Barcodes were selected from barcodes used in NEBNext® Multiplex Oligos for Illumina® (E73325, NEB). The resulting set was 96 forward x 16 reverse barcodes enabling the multiplexed sequencing of 1536 isolates in a single pool.

Colonies were picked into 10 μL of 25% glycerol and 50% LB mixture in water. 0.8 μL of this was then used to template an 8 μL PCR using LongAmp® Hot Start *Taq* 2X Master Mix (M0533, NEB) with each amplification containing a unique combination of forward and reverse sequencing-barcoded primers that were cherry-picked into PCR reactions using Echo 525 with a 384PP plate as was done in the construction. PCR reactions were first run at 94 °C for 1 min followed by 35 cycles of 94 °C for 15 seconds, 53 °C for 15 seconds, and 65 °C for 3 minutes. The final hold was for 10 minutes at °C.

The resulting amplicons were pooled in equal volumes and cleaned up using 1.8X SPRISelect beads. DNA was eluted in nuclease free water and DNA concentration was measured using a NanoDrop™. 1.2 μg of DNA was then used as input for a Ligation Sequencing Kit (SQK-LSK109, Oxford Nanopore, Oxford, United Kingdom) with the NEBNext® Companion Module for Oxford Nanopore Technologies® Ligation Sequencing (E7180, NEB). The Oxford Nanopore protocol, Amplicons by Ligation (SQK-LSK109) Version: ACDE_9064_v109_revP_14Aug2019 was followed. Data was acquired for 24-72 hours.

After sequencing, all reads were demultiplexed using minibar software^24^ (https://github.com/calacademy-research/minibar) based on the specified combination of sequences of barcodes and primers at the 5’ and 3’ ends for each isolate. The predominant consensus sequence was derived for each isolate using Amplicon_sorter software^25^.

Sequencing results were then confirmed by mapping a library of digested fragment annotations to the consensus sequences using Geneious (Biomatters, Inc. Boston, MA USA). Fragments were manually scored as either correct (matching the expected sequence), containing all parts (all fragments were properly assembled with at least 10 bp from each fragment), missing pieces (not all fragments were present), wrong fragments (fragments from other assemblies), vector closure (either parent vector or closure without inserts), or no amplicon (colony PCR failed to produce an amplicon for sequencing).

## Acknowledgements

We would like to thank Peter Weigele, Yan-Jiun Lee, Mehmet Berkmen, James Eaglesham, Emily McNutt, Lana Saleh, Katherine O’Toole, Andy Gardner, and Chen Song for providing sequences in which to validate these methods. Funding for open access is from New England Biolabs.

## Online Supplement

**Supplementary Figure 1.**
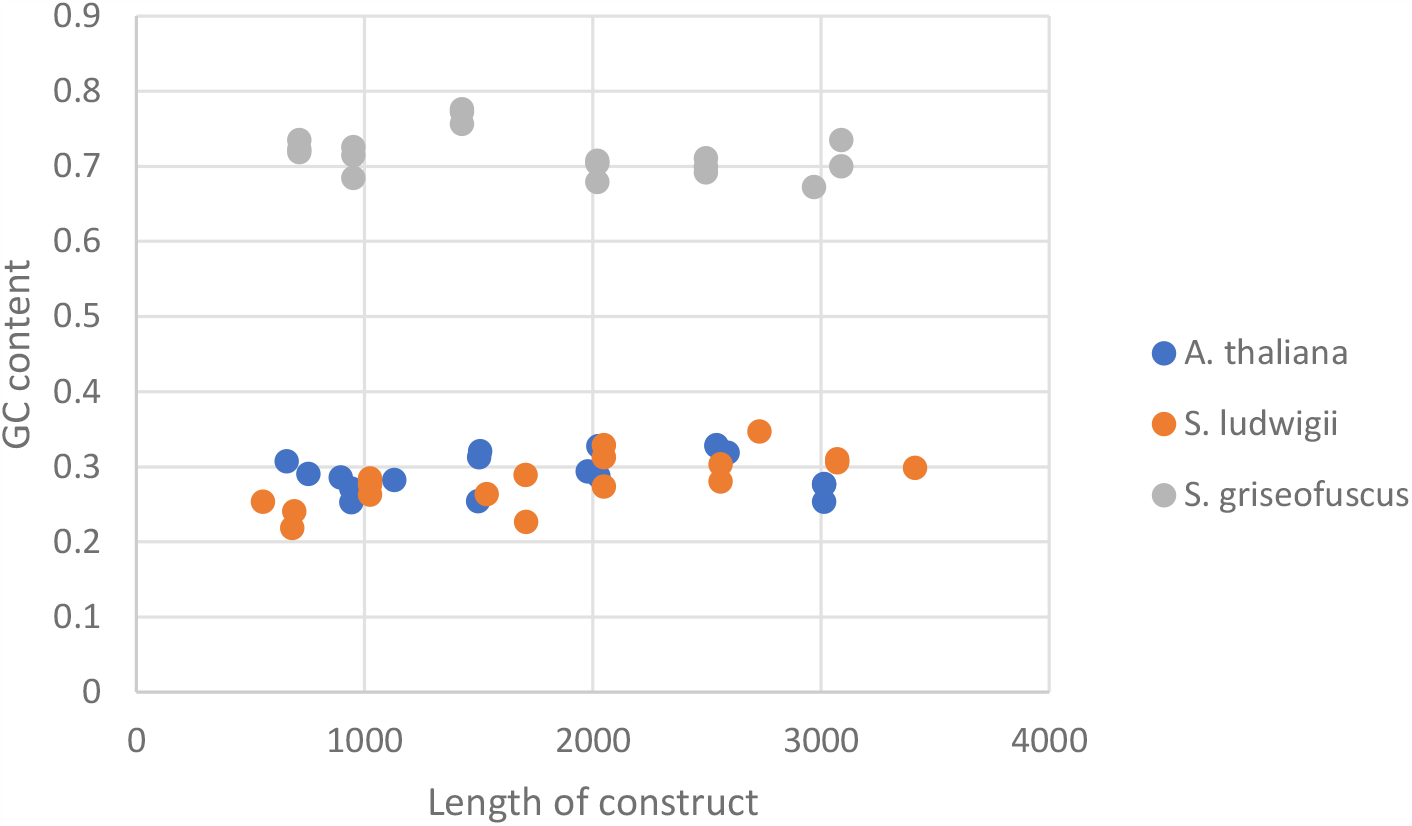
GC content of challenging sequences constructed.

**Supplementary Figure 2.**
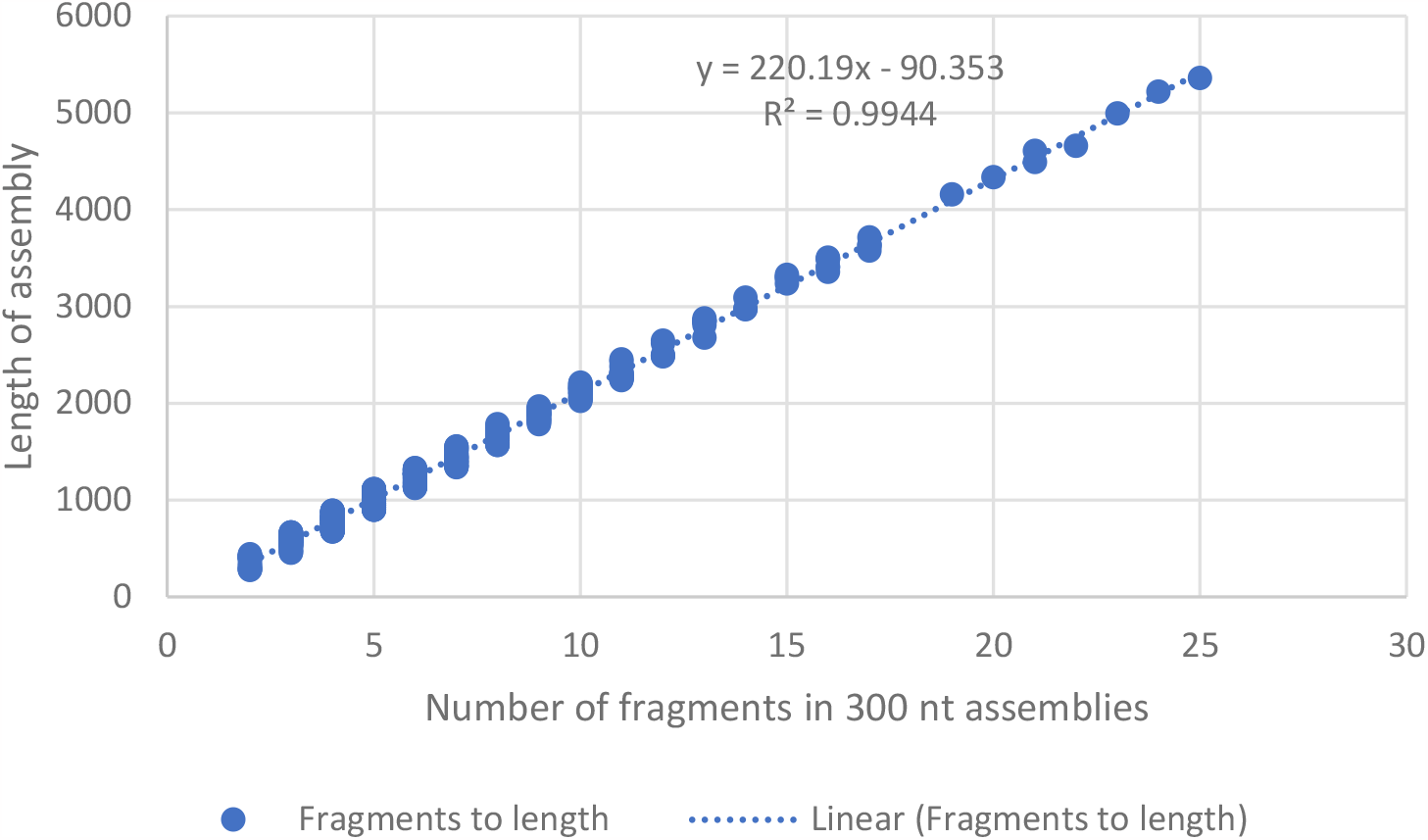
Correlation between number of fragments in 300 nt oligo pool to length of entire construct. Each fragment represents approximately 220 bp of DNA construct.

**Supplementary Figure 3.**
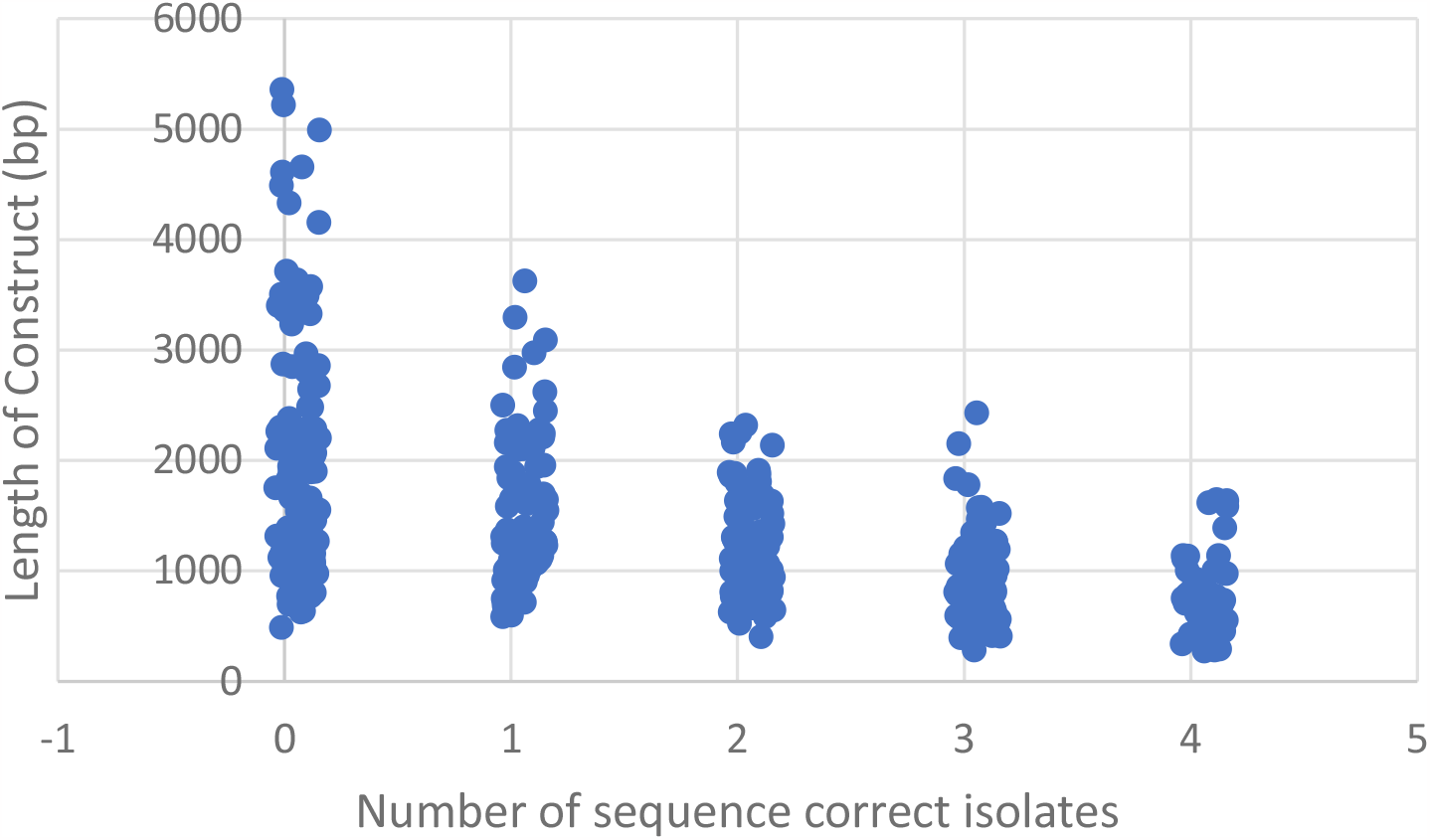
Correlation between the number of sequence correct colonies and length of construct. Number of sequence correct colonies discrete but plotting is jittered to improve visualization.

**Supplementary Figure 4.**
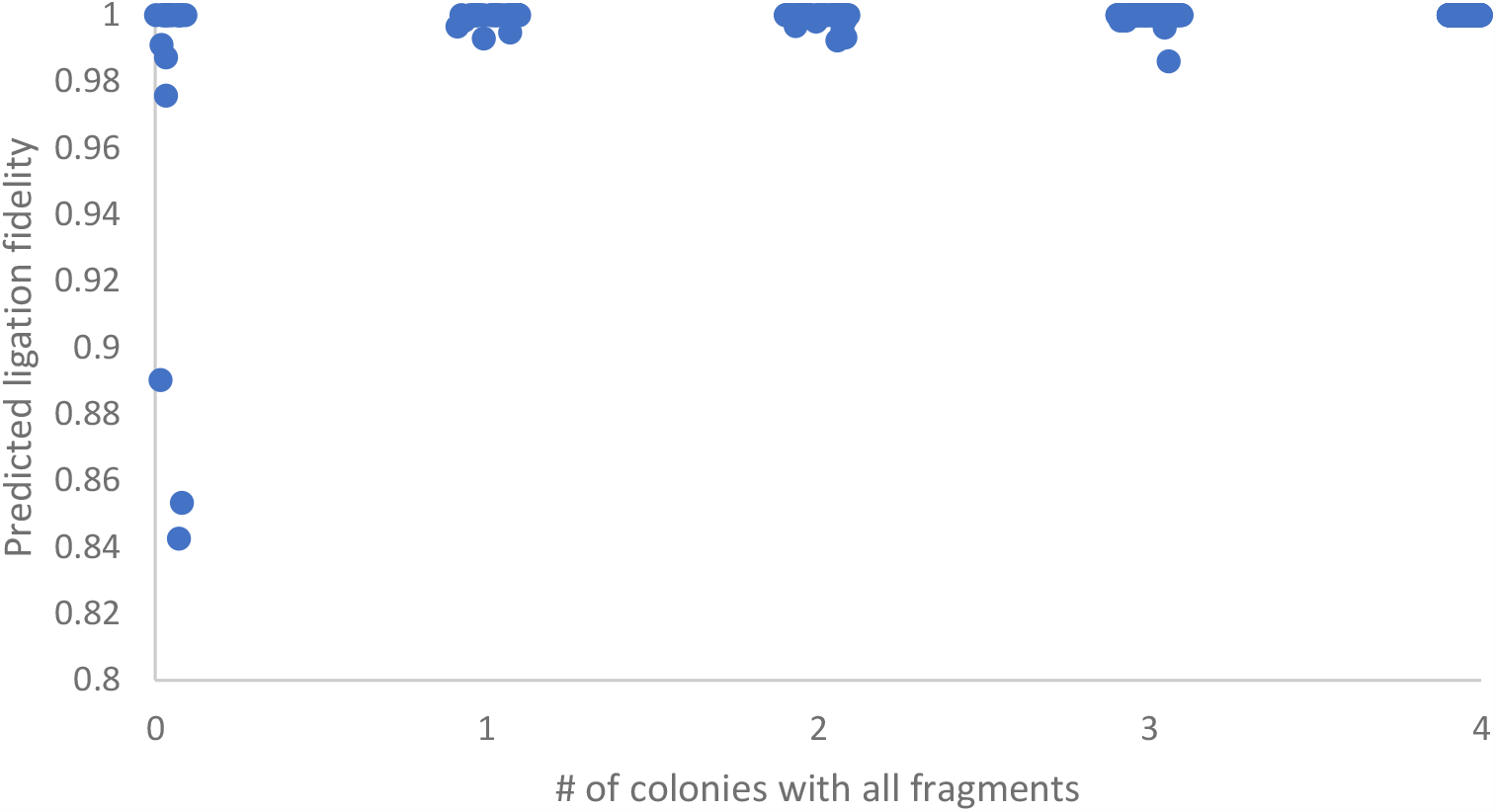
Correlation between predicted ligation fidelity and the number of colonies with all fragments present. Number of colonies with all fragments is discrete but plotting is jittered to improve visualization.

**Supplementary Figure 5.**
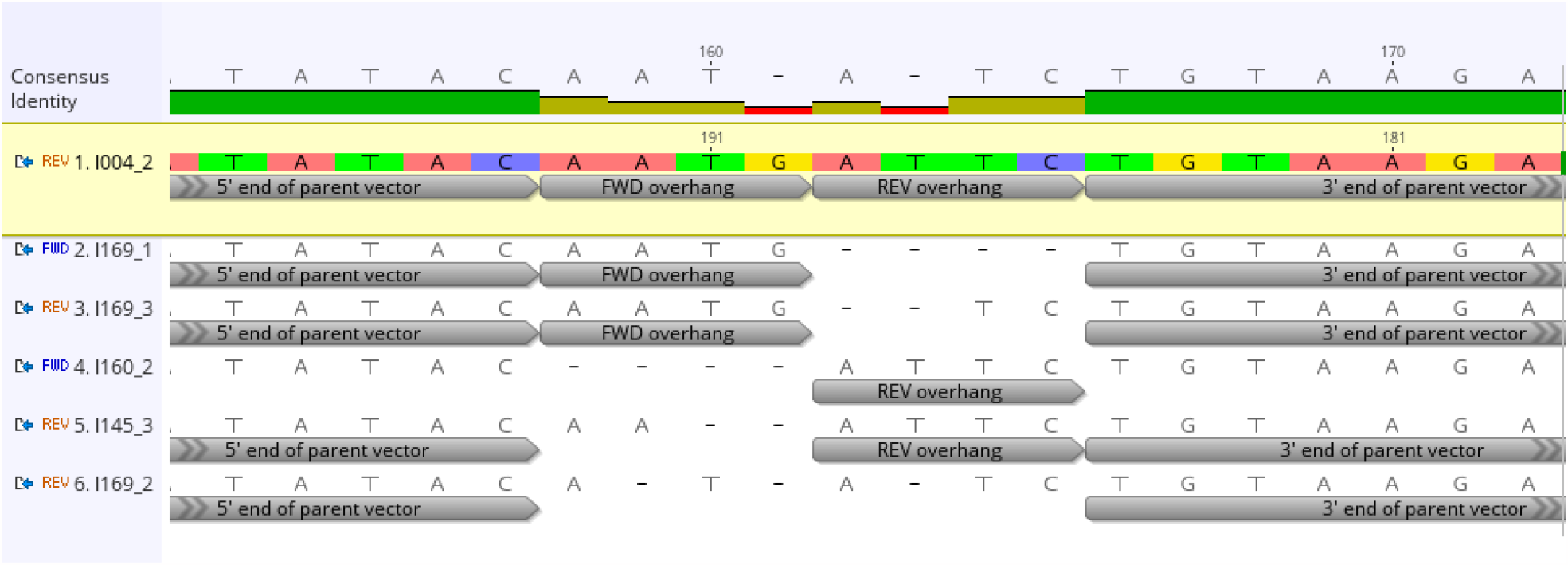
Failure mechanisms of parental vector closure. The top line represents the consensus sequence of the predominant failure mechanism resulting from digestion of the multiple cloning site followed by 5’-3’ end filling and blunt-end ligation.

**Supplementary Table 1.**
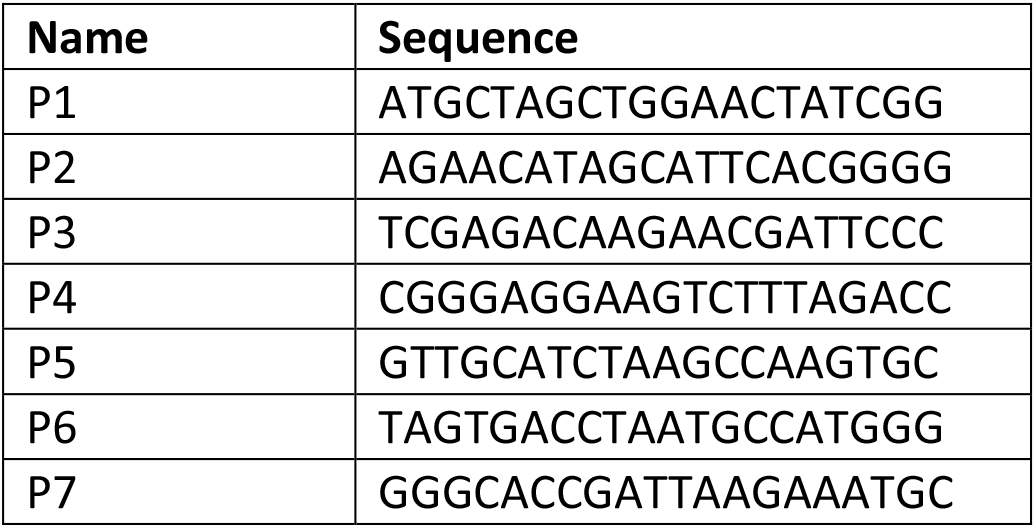
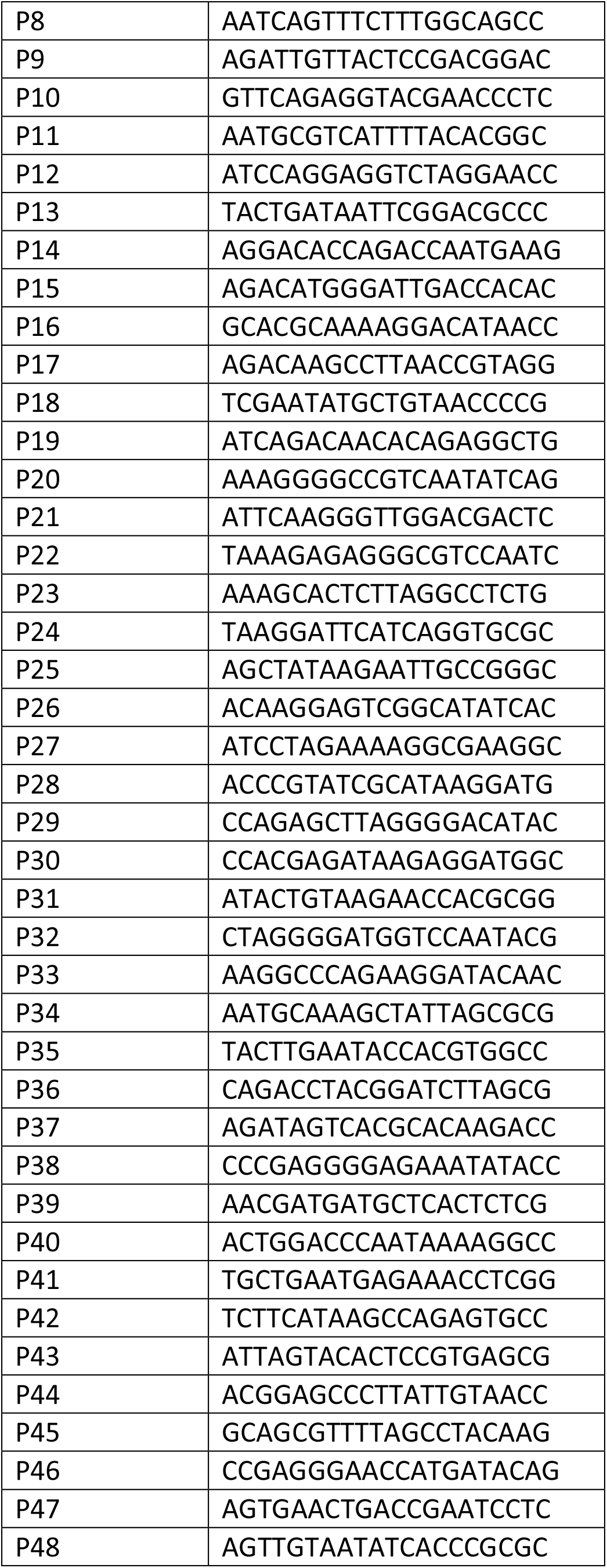
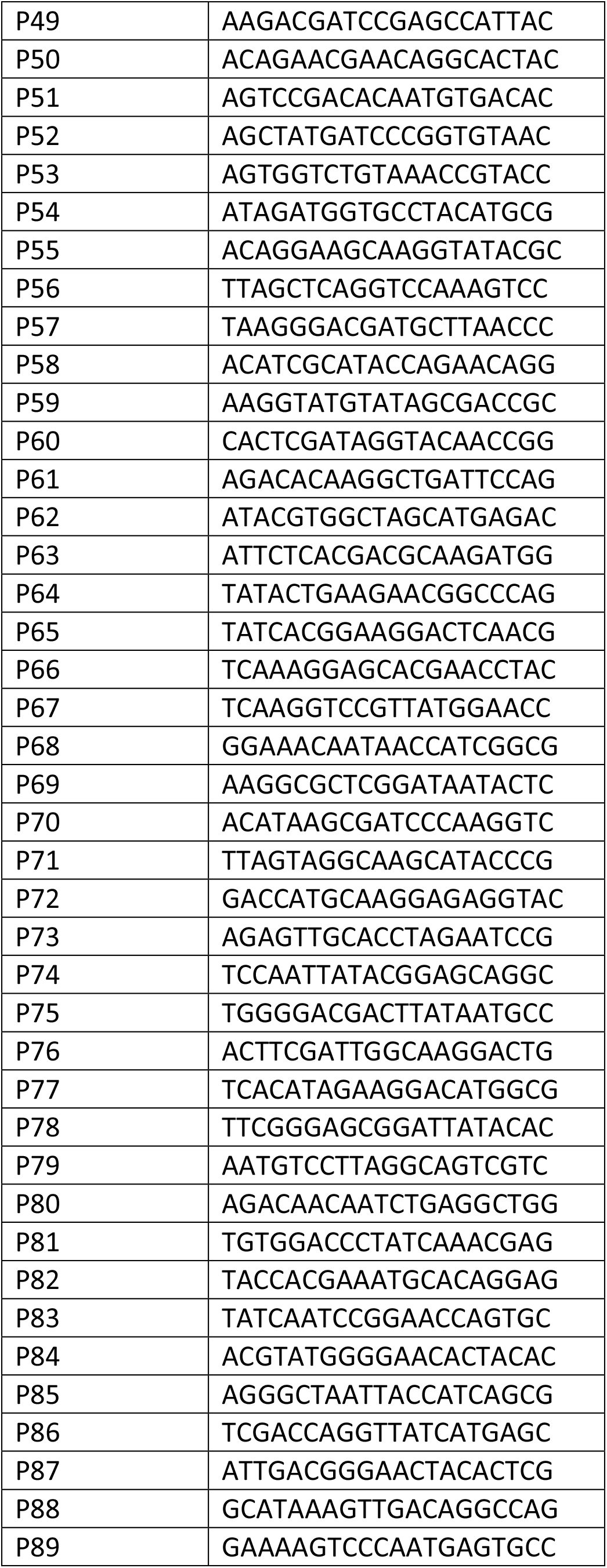
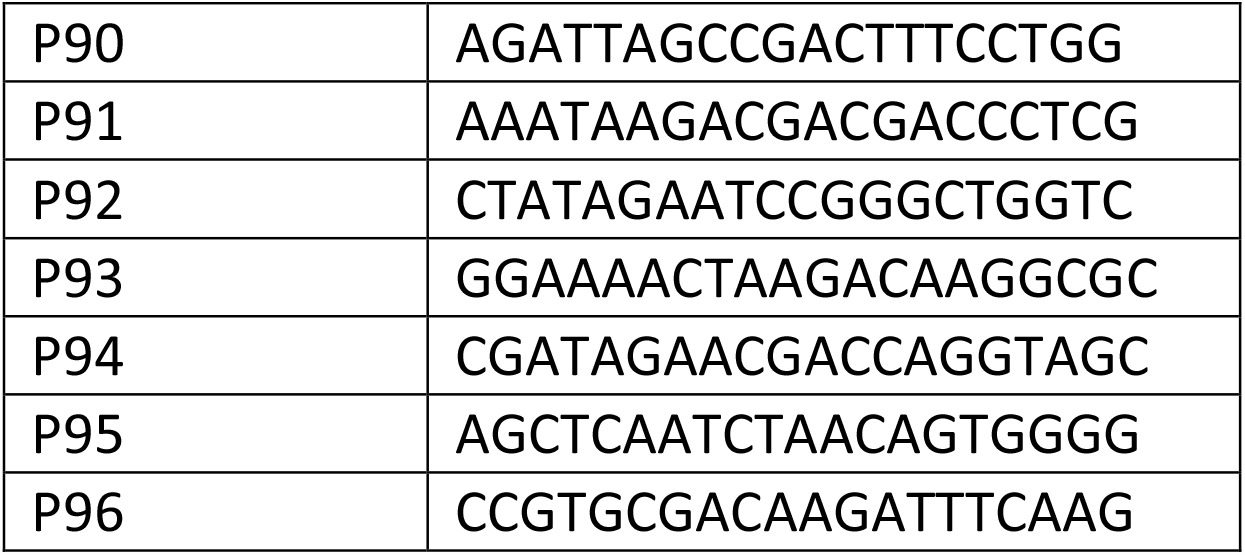
Barcodes used for sequence retrieval.

**Supplementary File 1**. Synthesis of difficult sequences using different layouts.

**Supplementary File 2**. Synthesis of 458 genes.

**Supplementary File 3**. Genbank file of constitutive vector for GGA.

**Supplementary File 4**. Genbank file of domesticated pET vector for GGA.

